# NanoCLUST: a species-level analysis of 16S rRNA nanopore sequencing data

**DOI:** 10.1101/2020.05.14.087353

**Authors:** Héctor Rodríguez-Pérez, Laura Ciuffreda, Carlos Flores

**Affiliations:** Research Unit, Hospital Universitario N.S. de Candelaria, Universidad de La Laguna, 38010, Santa Cruz de Tenerife, Spain; CIBER de Enfermedades Respiratorias, Instituto de Salud Carlos III, 28029, Madrid, Spain; Genomics Division, Instituto Tecnológico y de Energías Renovables (ITER), 38600 Granadilla, Santa Cruz de Tenerife, Spain; Instituto de Tecnologías Biomédicas (ITB), Universidad de La Laguna, 38200 San Cristóbal de La Laguna, Santa Cruz de Tenerife, Spain

## Abstract

**Summary:** NanoCLUST is an analysis pipeline for classification of amplicon-based full-length 16S rRNA nanopore reads. It is characterized by an unsupervised read clustering step, based on Uniform Manifold Approximation and Projection (UMAP), followed by the construction of a polished read and subsequent Blast classification. Here we demonstrate that NanoCLUST performs better than other state-of-the-art software in the characterization of two commercial mock communities, enabling accurate bacterial identification and abundance profile estimation at species level resolution.

**Availability and implementation:** Source code, test data and documentation of NanoCLUST is freely available at https://github.com/genomicsITER/NanoCLUST under MIT License.

## 1 Introduction

Nanopore sequencing (Oxford Nanopore Technologies, ONT) has emerged as a fast and inexpensive method for long-read DNA/RNA sequencing. Accessing the microbial communities with ONT is feasible using rapid protocols targeting the full-length 16S rRNA gene. Widely used software, such as QIIME (Caporaso *et al.*, 2010), is designed to analyze short-read sequencing that typically allow characterizing the microbial communities at the genus level. However, tools for the analysis of the noisy 16S rRNA long-reads from ONT are scarce (Santos *et al.*, 2020). Among them, the popular Epi2me (https://www.metrichor.com) is based on a read-by-read classification strategy that does not cope well with the error rate associated to this technology, resulting in the misclassification of a high percentage of reads and a high uncertainty in the results.

Here we present NanoCLUST, a pipeline for the analysis of ONT 16S rRNA amplicon reads. Besides demultiplexing and quality control (QC) steps, and inspired by the work of Beaulaurier *et al.* (2020), it leverages Uniform Manifold Approximation and Projection (UMAP) (McInnes *et al.*, 2018) and Hierarchical Density-Based Spatial Clustering of Applications with Noise (HDBSCAN) (McInnes *et al.*, 2017) for unsupervised read clustering followed by the construction of a polished sequence for subsequent taxonomic classification. We tested NanoCLUST on ONT data from two commercial mock communities and compared the results to two other popular and accurate classification methods, Kraken2 (Wood *et al.*, 2019) and Bracken (Lu *et al.*, 2017).

## 2 Description and implementation

NanoCLUST is implemented in Nextflow workflow management system, which enables efficient parallel execution in all major systems and computing environments. NanoCLUST development followed the nf-core (Ewels *et al.*, 2020) best practices guidelines and standardized template. Software packages and the dependencies are bundled in the pipeline using built-in integration for conda environments and Docker containers. The general workflow of NanoCLUST is illustrated in Figure 1, and software versions detailed in Table S1. The input data consists of basecalled 16S rRNA ONT sequencing reads, which are internally demultiplexed in case of pooled samples. Then, reads are filtered using fastp to ensure that only near full-length 16S rRNA reads are kept. By default, reads with a Phred score <8.0 and of length below 1,400 base pairs (bp) or above 1,700 bp are discarded. Prior to the clustering step, each read is transformed into a normalized 5-mer frequency vector and stored in tabular format. UMAP projection and posterior clustering using HDBSCAN is applied to the 5-mer frequency vector set and a cluster assignation is given to each read. The clustering step parameters, such as the minimum read number necessary to identify an independent cluster to determine the sensitivity, as well as other HDBSCAN parameters can be manually set by the user. These parameters will be dependent on the input data and the desired level of taxonomy. Therefore, they have a significant practical effect on the clustering and on subsequent analysis.

**Figure 1.**
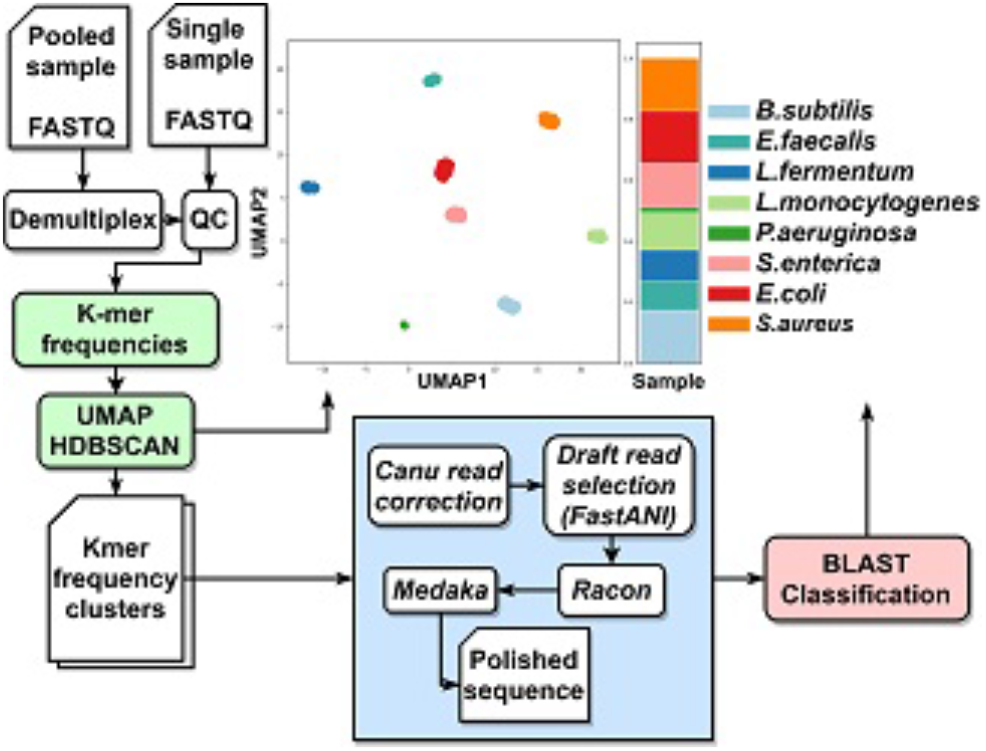
Simplified flowchart of NanoCLUST.

The next step builds a consensus sequence from the reads belonging to each cluster. For that, the pairwise Average Nucleotide Identity (ANI) between reads in the same cluster is calculated using FastANI. Then, the read with the highest average intra-cluster ANI is chosen and 100 other reads from the same cluster are selected for polishing the sequence. The polishing stage includes one round each in the Canu read-correction module, in Racon, and in Medaka. The resulting polished sequence is taken as the representative of the cluster and is finally classified using blastn and the NCBI Refseq database (or any other database provided by the user). The resulting taxonomic classification output is assigned to all the reads belonging to the cluster and the results are then merged for relative abundance estimation based on the HDBSCAN cluster assignation and the total number of reads projected by UMAP. The output consists of QC reports of the input data, the consensus sequence representing each cluster, a complete classification report for each cluster, and the relative abundance tables and bar plots at multiple taxonomic levels, both for single and pooled samples.

## 3 Results

We used NanoCLUST to analyze the sequencing data from two commercial mock communities run in various experiments, MOCK1 containing genomic DNA from eight bacterial species (Zymobiomics), and MOCK2 containing genomic DNA from 20 bacterial species (Bioresources) (see Supplementary data). Eight clusters were identified in MOCK1, and classification of the polished sequences successfully detected all taxa present in the sample at the species level (Figure S1). As regards of MOCK2, 19 out of the 20 expected clusters were identified (Figure S2). Of these, only one species, *Bacillus cereus*, was misclassified as *Bacillus thuringiensis*, likely because of the 99.73% similarity in their 16S rRNA sequence (Figure S3). The absence of one species, corresponding to *Actinomices odontolyticus*, was explained by a low amplicon representation in the sequencing experiment, not reaching the minimum read number set (n=100) for UMAP cluster formation.

We then compared the results obtained from NanoCLUST for the two communities with those from Kraken2 (Wood *et al.*, 2019) and Bracken (Lu *et al.*, 2017) (Figure S4, S5). Overall, NanoCLUST performed better than the other classifiers in the analysis of 16S rRNA ONT sequences. Species richness estimated by NanoCLUST in the independent sequencing runs was the most similar to the expected, with eight and 19 species identified for MOCK1 and MOCK2, respectively, while Kraken2 and Bracken identified a much larger number of species in the two communities (an average of 139 and 214 different species for MOCK1 and MOCK2, respectively). Shannon diversity index was also calculated, and the closer value to the expected for each mock was found when NanoCLUST was used in the analysis, compared to Kraken2 and Bracken (Figure S6).

To compare the expected relative abundance of species in the two mock communities with the estimates provided by the three classification methods, we calculated the mean absolute error (MSA) and the root mean squared error (RMSE). As indicated by the significantly lower MSA and RMSE, relative abundances obtained by NanoCLUST were more similar to the expected than those of Kraken2 and Bracken (Figure S7, S8).

## Acknowledgements

We would like to thank Tamara Hernandez-Beeftink and José M. Lorenzo-Salazar for their help with the experimental setup with the MinION device and the practical advice with UMAP/HDBSCAN parameters, respectively.

## Funding

This work was supported by Instituto de Salud Carlos III [PI14/00844, PI17/00610, and FI18/00230] and co-financed by the European Regional Development Funds, “A way of making Europe” from the European Union; Ministerio de Ciencia e Innovación [RTC-2017-6471-1, AEI/FEDER, UE]; Cabildo Insular de Tenerife [CGIEU0000219140]; Fundación Canaria Instituto de Investigación Sanitaria de Canarias [PIFUN48/18]; and by the agreement with Instituto Tecnológico y de Energías Renovables (ITER) to strengthen scientific and technological education, training, research, development and innovation in Genomics, Personalized Medicine and Biotechnology [OA17/008].

## Conflict of Interest

none declared.

## Supplementary information

**Table S1.** Software and algorithms integrated in NanoCLUST*.

| Software/algorithm | Utility | Version |
| --- | --- | --- |
| qcat<sup>a</sup> | Demultiplexing | 1.1.0 |
| Porechop<sup>b</sup> | Demultiplexing | 0.2.3 |
| fastp<sup>c</sup> | QC filtering | 0.20.0 |
| fastQC<sup>d</sup> | QC report | 0.11.8 |
| multiQC<sup>e</sup> | QC report | 1.6 |
| UMAP<sup>f</sup> | Read projection | 0.3.10 |
| HDBSCAN<sup>g</sup> | Cluster assignation | 0.8.26 |
| seqtk<sup>h</sup> | Cluster read extraction | 1.3 |
| FastANI<sup>i</sup> | Draft selection | 1.3 |
| Canu<sup>j</sup> | Read correction | 2.0 |
| Racon<sup>k</sup> | Read polishing | 1.4.13 |
| Medaka<sup>l</sup> | Read polishing | 0.11.5 |
| blastn<sup>m</sup> | Taxonomic assignation | 2.9.0 |

<sup>a</sup>https://github.com/nanoporetech/qcat; <sup>b</sup>https://github.com/rrwick/Porechop;
<sup>c</sup>Chen *et al.* (2018); <sup>d</sup>Andrews S. (2010); <sup>e</sup>Ewels *et al.* (2016); <sup>f</sup>McInnes *et al.* (2018); <sup>g</sup>McInnes *et al.* (2017); <sup>h</sup>https://github.com/lh3/seqtk; <sup>i</sup>Jain *et al.* (2018); <sup>j</sup>Koren *et al.* (2017); <sup>k</sup>Vaser *et al.* (2017); <sup>l</sup>https://github.com/nanoporetech/medaka; <sup>m</sup>Altschul *et al.* (1990). \*Includes the following python libraries for data processing and plotting: Python 3.7, Numpy 1.18.1, Pandas 1.0.3, and Matplotlib 3.2.1.

### Library preparation and sequencing

Amplicon-based libraries were prepared using the ONT 16S Barcoding kit (SQK-RAB204) following manufacturer’s instructions (ONT). Briefly, 10 ng of DNA from two bacterial mock communities (ZymoBIOMICS™ Microbial Community DNA Standard [Zymo Research]; Microbial Mock Community B (Even, High Concentration) [BEI Resources]) were used as targets in the PCR amplification reaction, performed using the LongAmp Taq 2X MasterMix (New England Biolabs) in a final volume of 50 μl. Negative controls (containing only PCR grade water) were also included in each reaction. PCR products were purified using 30 μl of the AMPure XP beads (Beckman Coulter) and eluted in 10 μl of 10mM Tris-HCl pH 8.0 with 50 mM NaCl. Each barcoded library was quantified using the Invitrogen™ Qubit™ dsDNA HS Assay kit (Thermo Fisher Scientific) and pools of 12-plex barcoded libraries were prepared for each run. Rapid adapters (1 μl) were added to each library pool and followed by an incubation at 23 °C for 5 minutes. Libraries were sequenced using a MinION device (ONT) over a period of 48 h. After priming and loading of the flow cell (R9.4.1), the run was started using MinKNOW software (v19.05). Fast5 files were generated and basecalled using Guppy (v3.1.5) using a local machine with two Xeon 6238T 1,9 GHz, GPU Nvidia RTX 2080TI and 512GB RAM. Demultiplexing and quality controls were carried out either using the NanoCLUST modules for this purpose, or local qcat Python package (ONT) and fastp (v0.20.0) (Chen *et al.*, 2018).

### Comparative analysis of relative abundances

Seven and three sequencing replicates were carried out using the ZymoBIOMICS Microbial DNA Standard (MOCK1) and the Microbial MOCK Community B (MOCK2), respectively. A subset of 100,000 high-quality reads (mean length > 1,400 bp and < 1,700 bp and a Phred quality score > 8.0) from each mock replicate was analyzed using NanoCLUST. The same subset of reads was subsequently analyzed using Kraken2 (v2.0.8-beta) (Wood *et al.*, 2019) and Bracken (v2.5.0) (Lu *et al.*, 2017) against the Refseq complete bacterial genomes database. This subset (only from MOCK1) is included in the repository as an example test file. Bacterial abundance profiles were retrieved from the Kraken2 and Bracken reports and compared to those calculated by NanoCLUST. Species richness and Shannon diversity index were retrieved using the vegan R package (Oksanen *et al.*, 2019). The metrics to assess the deviations from the expected values (the mean absolute error (MAE) and the root mean squared error (RMSE)) were calculated using R software (R Core Team, 2013), and mean values compared between groups using ANOVA followed by the Tukey’s multiple comparison test.

### Supplementary figures

**Figure S1.**
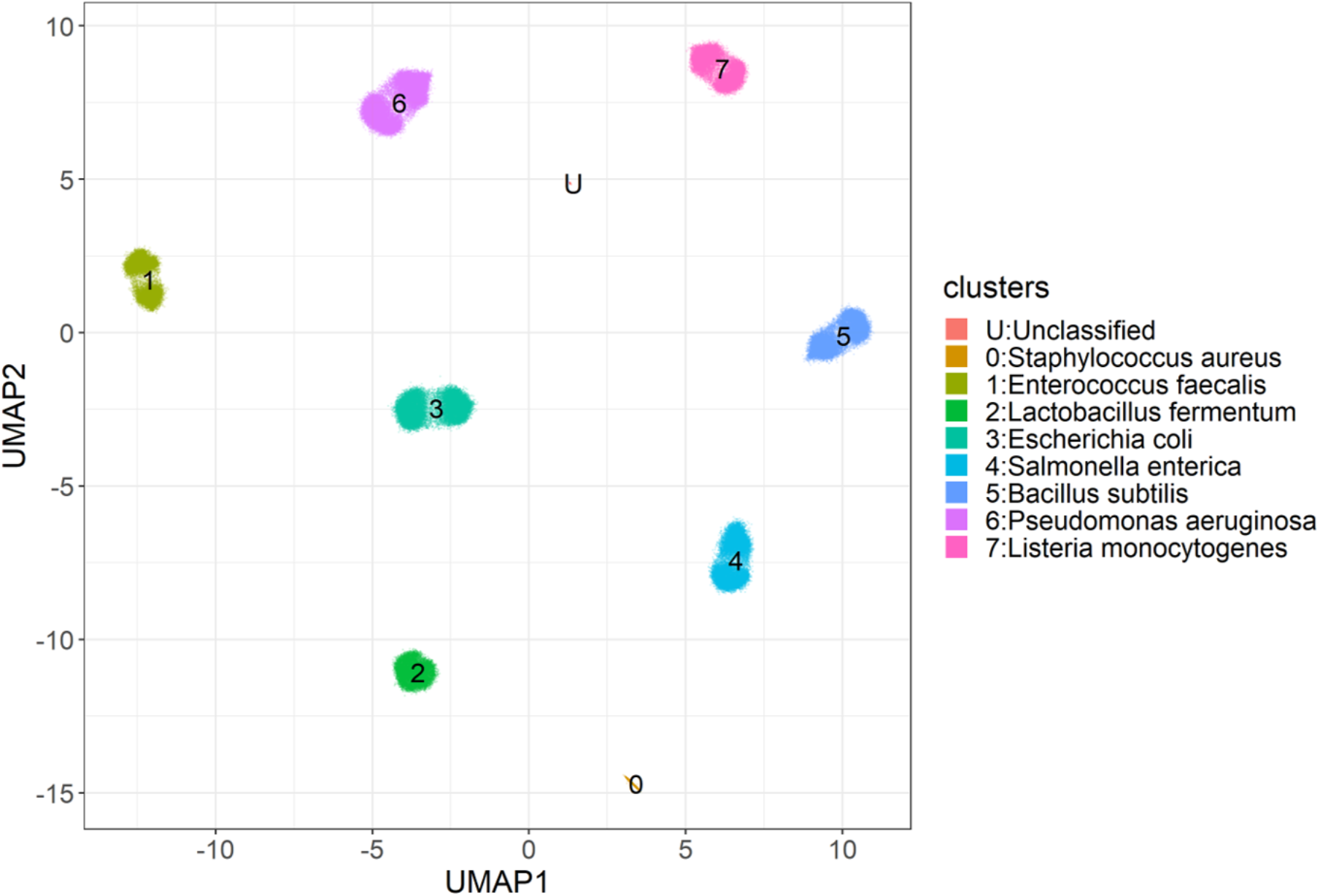
UMAP plot of reads from MOCK1.

**Figure S2.**
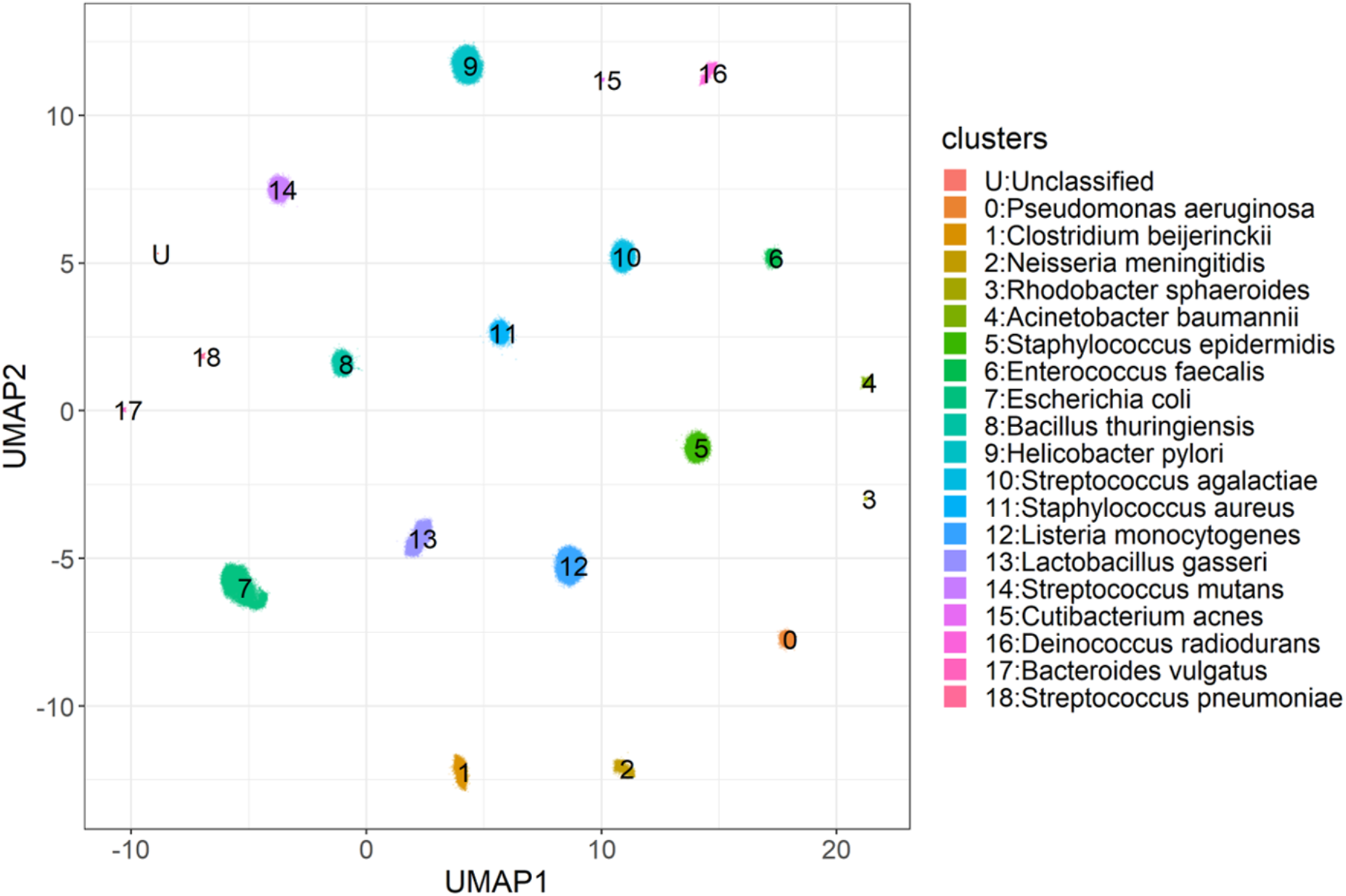
UMAP plot of reads from MOCK2. Note that *Bacillus thuringiensis*, supported by NanoCLUST, is not present in this mock community.

**Figure S3.**
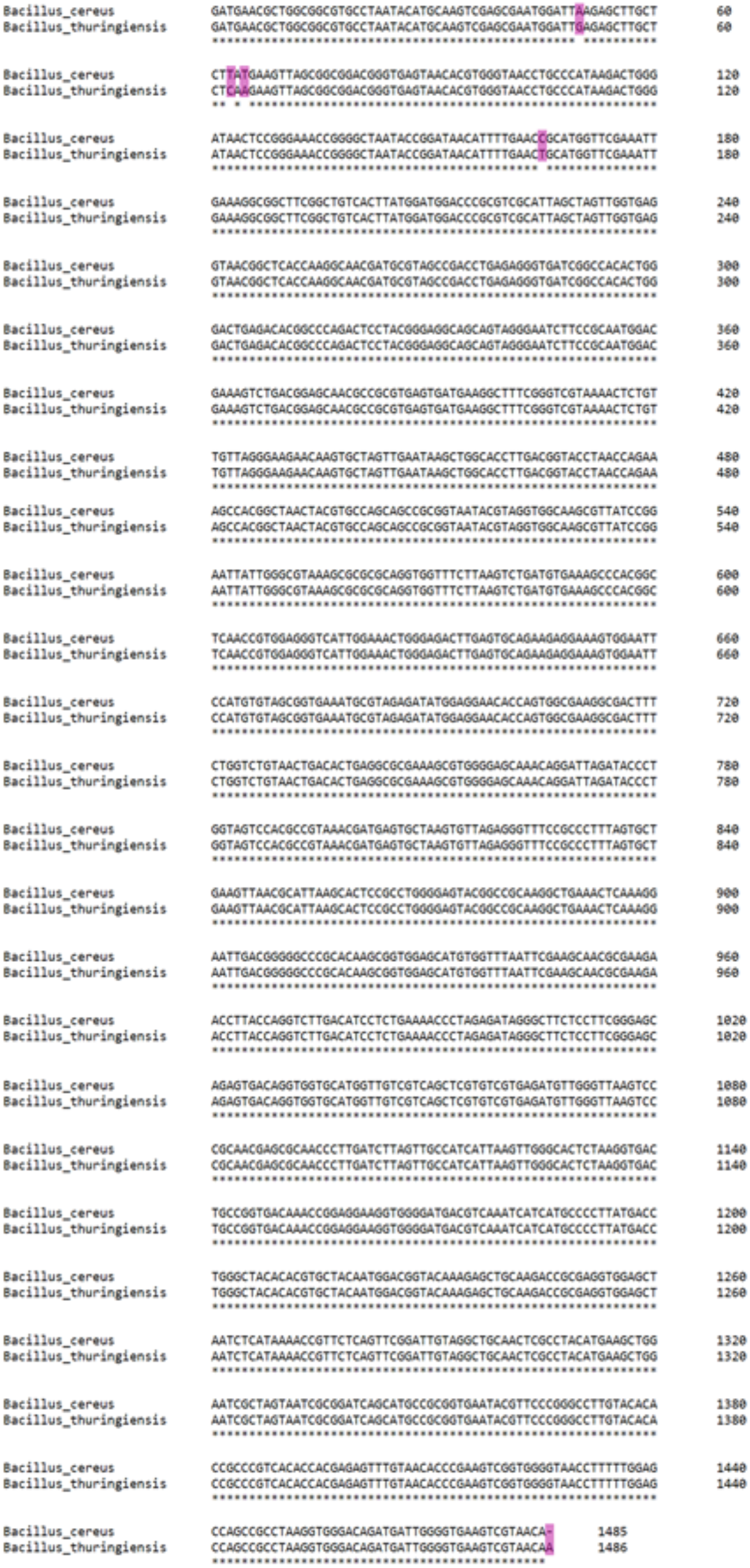
Sequence alignment (mismatches in purple) of 16S rRNA genes from *Bacillus cereus* and *Bacillus thuringiensis*. Alignment was performed using Clustal Omega (Sievers *et al.*, 2011).

**Figure S4.**
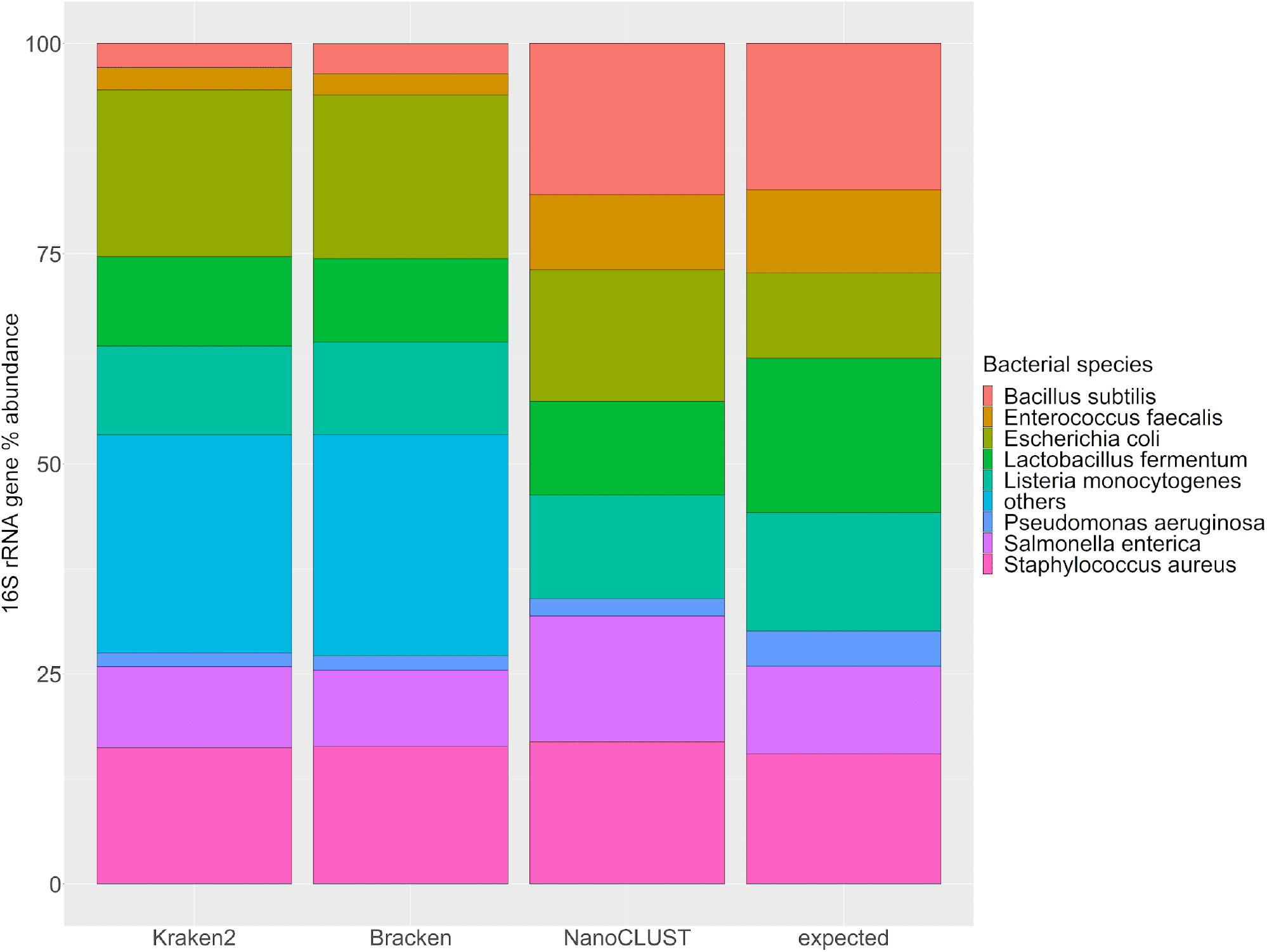
Relative abundances of Kraken2, Bracken, and NanoCLUST compared to the expected values for MOCK1.

**Figure S5.**
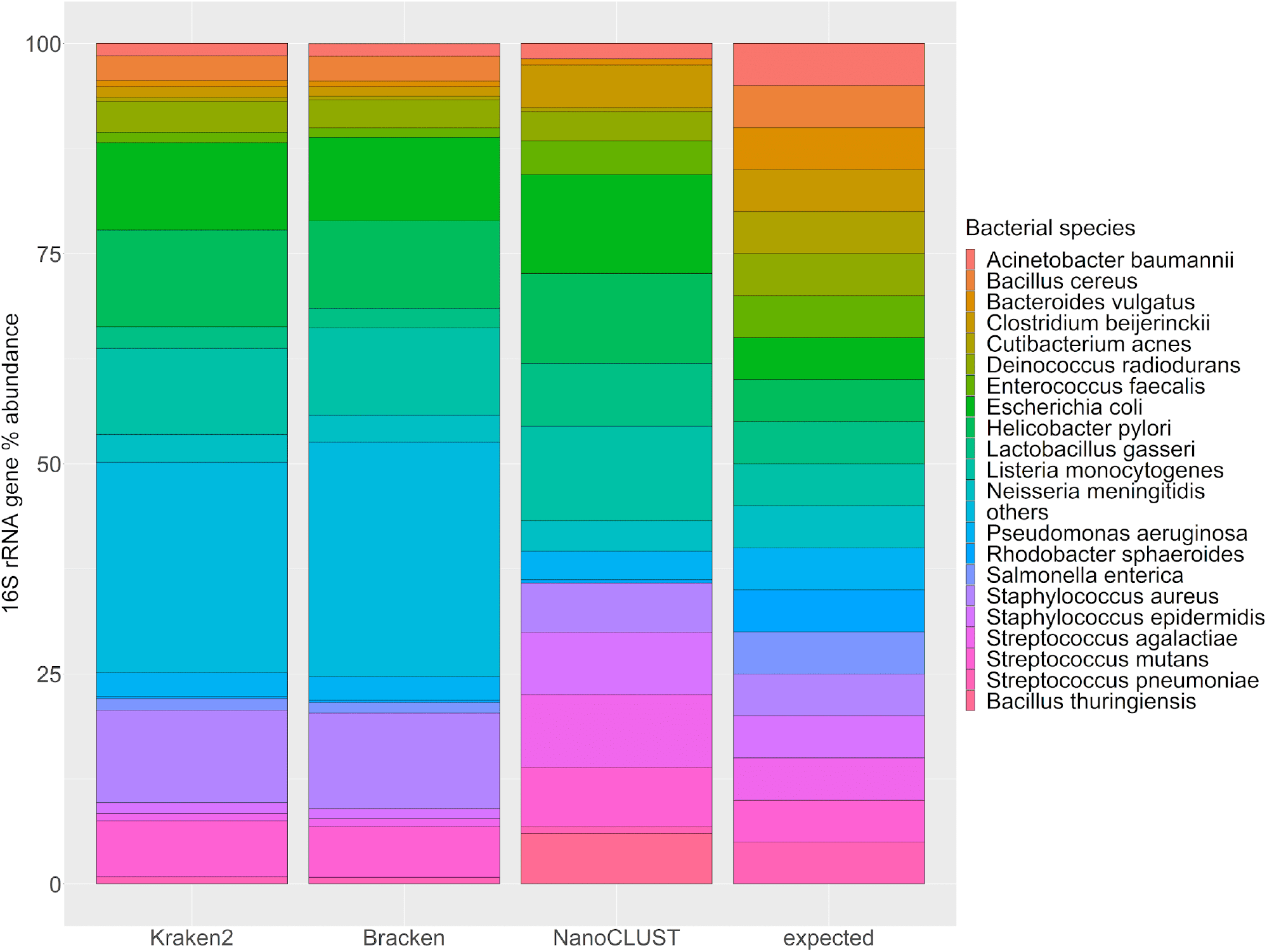
Relative abundances of Kraken2, Bracken, and NanoCLUST compared to the expected values for MOCK2. Note that *Bacillus thuringiensis*, supported by NanoCLUST, is not present in this mock community.

**Figure S6.**
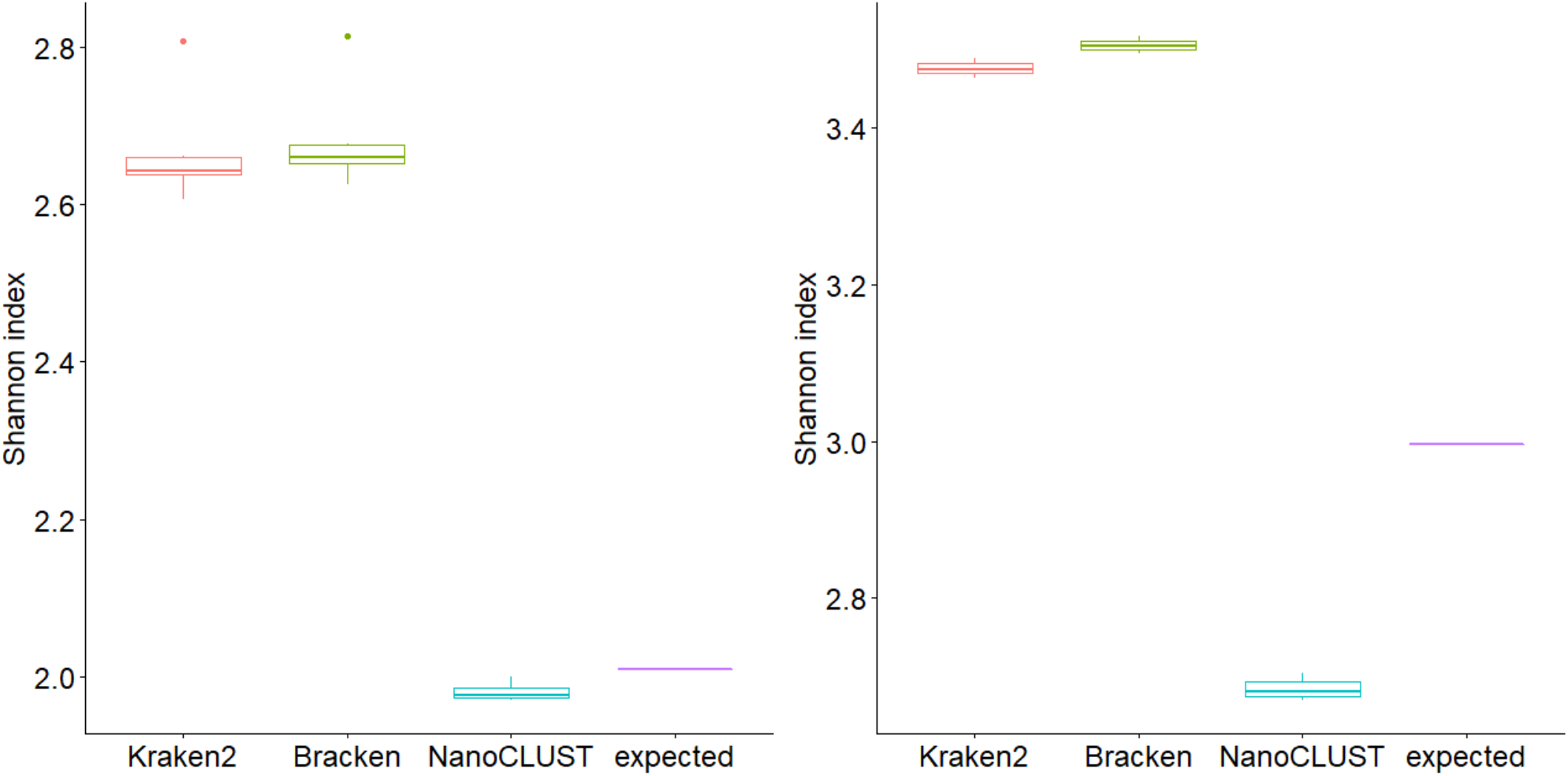
Shannon diversity index for MOCK1 (left panel) and MOCK2 (right panel).

**Figure S7.**
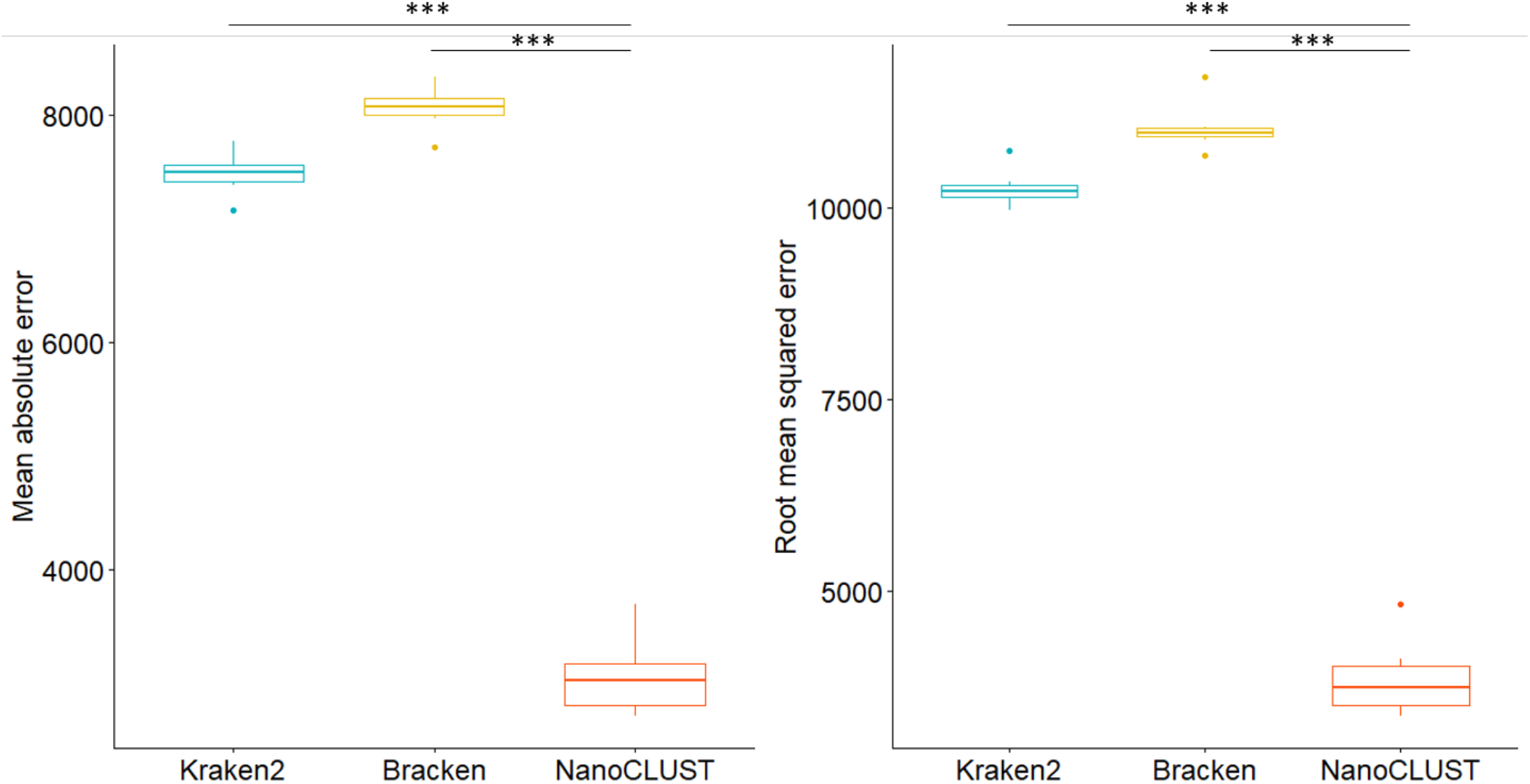
Mean absolute error and root mean squared error against the expected of MOCK1, based on read counts. Tukey multiple comparison test. ***, p<0.001.

**Figure S8.**
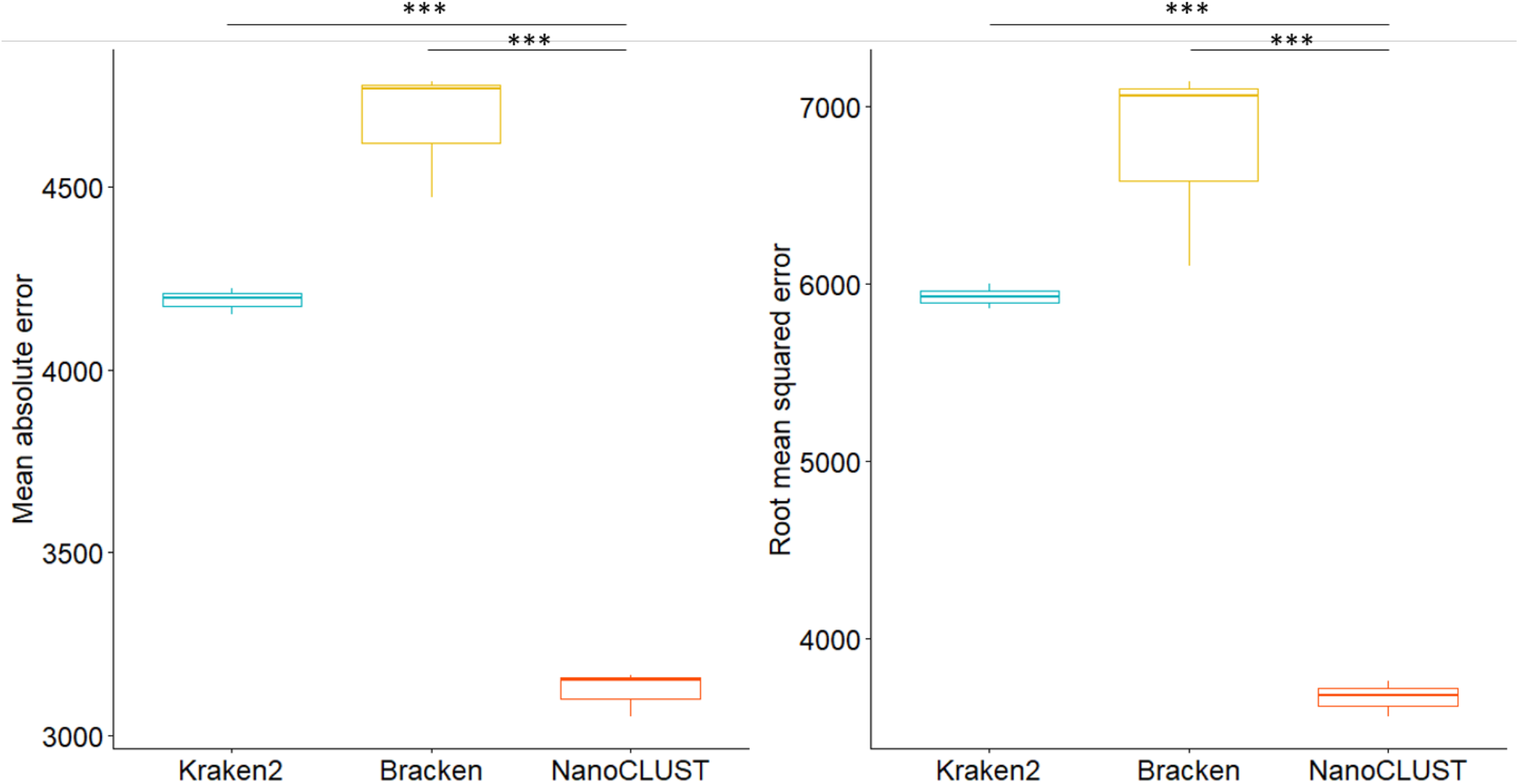
Mean absolute error and root mean squared error against the expected of MOCK2, based on read counts. Tukey multiple comparison test. ***, p<0.001.

## Notes

https://github.com/genomicsITER/NanoCLUST

### Competing Interest Statement

The authors have declared no competing interest.

